# Indole-3-Acetaldehyde Dehydrogenase-dependent Auxin Synthesis Contributes to Virulence of *Pseudomonas syringae* Strain DC3000

**DOI:** 10.1101/173302

**Authors:** Sheri A. McClerklin, Soon Goo Lee, Ron Nwumeh, Joseph M. Jez, Barbara N. Kunkel

**Affiliations:** Department of Biology, Washington University, St. Louis, Mo USA

## Abstract

The bacterial pathogen *Pseudomonas syringae* modulates plant hormone signaling to promote infection and disease development. *P. syringae* uses several strategies to manipulate auxin physiology in *Arabidopsis thaliana* to promote pathogenesis, including synthesis of indole-3-acetic acid (IAA), the predominant form of auxin in plants, and production of virulence factors that alter auxin responses in the host; however, the role of pathogen-derived auxin in *P. syringae* pathogenesis is not well understood. Here we demonstrate that *P. syringae* strain DC3000 produces IAA via a previously uncharacterized pathway and identify a novel indole-3-acetaldehyde dehydrogenase, AldA, that functions in IAA biosynthesis by catalyzing the NAD-dependent formation of IAA from indole-3-acetaldehyde (IAAld). Biochemical analysis and solving of the 1.9 Å resolution x-ray crystal structure reveal key features of AldA for IAA synthesis, including the molecular basis of substrate specificity. Disruption of *aldA* and a close homolog, *aldB*, lead to reduced IAA production in culture and reduced virulence on *A. thaliana.* We use these mutants to explore the mechanism by which pathogen-derived auxin contributes to virulence and show that IAA produced by DC3000 suppresses salicylic acid-mediated defenses in *A. thaliana.* Thus, auxin is a DC3000 virulence factor that promotes pathogenicity by suppressing host defenses.

**Author Summary:** Pathogens have evolved multiple strategies for suppressing host defenses and modulating host physiology to promote colonization and disease development. For example, the plant pathogen *Pseudomonas syringae* uses several strategies to the manipulate hormone signaling of its hosts, including production of virulence factors that alter hormone responses in and synthesis of plant hormones or hormone mimics. Synthesis of indole-3-acetic acid (IAA), a common form of the plant hormone auxin, by many plant pathogens has been implicated in virulence. However, the role of pathogen-derived IAA during pathogenesis by leaf spotting pathogens such as *P. syringae* strain DC3000 is not well understood. Here, we demonstrate that *P. syringae* strain DC3000 uses a previously uncharacterized biochemical pathway to synthesize IAA, catalyzed by a novel aldehyde dehydrogenase, AldA, and carry out biochemical and structural studies of the AldA protein to investigate AldA activity and substrate specificity. We also generate an *aldA* mutant disrupted in IAA synthesis to show that IAA is a DC3000 virulence factor that promotes pathogenesis by suppressing host defense responses.

## Introduction

Plant pathogens have evolved a variety of strategies to ensure a successful interaction with their hosts. These include the delivery of virulence proteins directly into host cells through the type III secretion system and production of plant hormones or hormone mimics. Both strategies are important for suppressing host defenses and/or modulating host physiology to promote colonization and disease development [1-3]. For example, the bacterial pathogen *Pseudomonas syringae*, the causal agent of bacterial speck disease [4, 5] produces the phytotoxin coronatine, a molecular mimic of the plant hormone jasmonic acid-isoleucine [6, 7]. Production and secretion of coronatine modulates host jasmonic acid signaling and is important for *P. syringae* pathogenesis [8-10]. Many plant-associated microbes also have the ability to synthesize indole-3-acetic acid (IAA), a common form of the phytohormone auxin, and in several cases production of IAA has been implicated in pathogen virulence [11, 12].

IAA synthesis in microbes has been well characterized, with five biosynthetic pathways for IAA utilizing the amino acid tryptophan (Trp) as the precursor (Fig 1) identified to date [11]. These include the indole-3-acetamide (IAM), the indole-3-acetonitrile (IAN), the indole-3-pyruvate (IPyA), the tryptamine (TAM), and the tryptophan side-chain oxidase (TSO) pathways [13]. Detailed analyses of the IAM and IPyA pathways helped elucidate the role of bacterial IAA production in plant-microbe interactions. Two enzymes responsible for converting Trp to IAA via the IAM pathway are tryptophan 2-monoxygenase (TMO) and IAM hydrolase (IAH), encoded by the *iaaM* and *iaaH* genes respectively [14]. Cloning of the *iaaM* and *iaaH* genes, as well as *ipdC* genes encoding IPyA decarboxylase [15-17], from a wide range of bacteria and the characterization of their encoded proteins provided insight on the various roles for IAA synthesis during pathogenesis [11, 18, 19].

**Fig 1.**
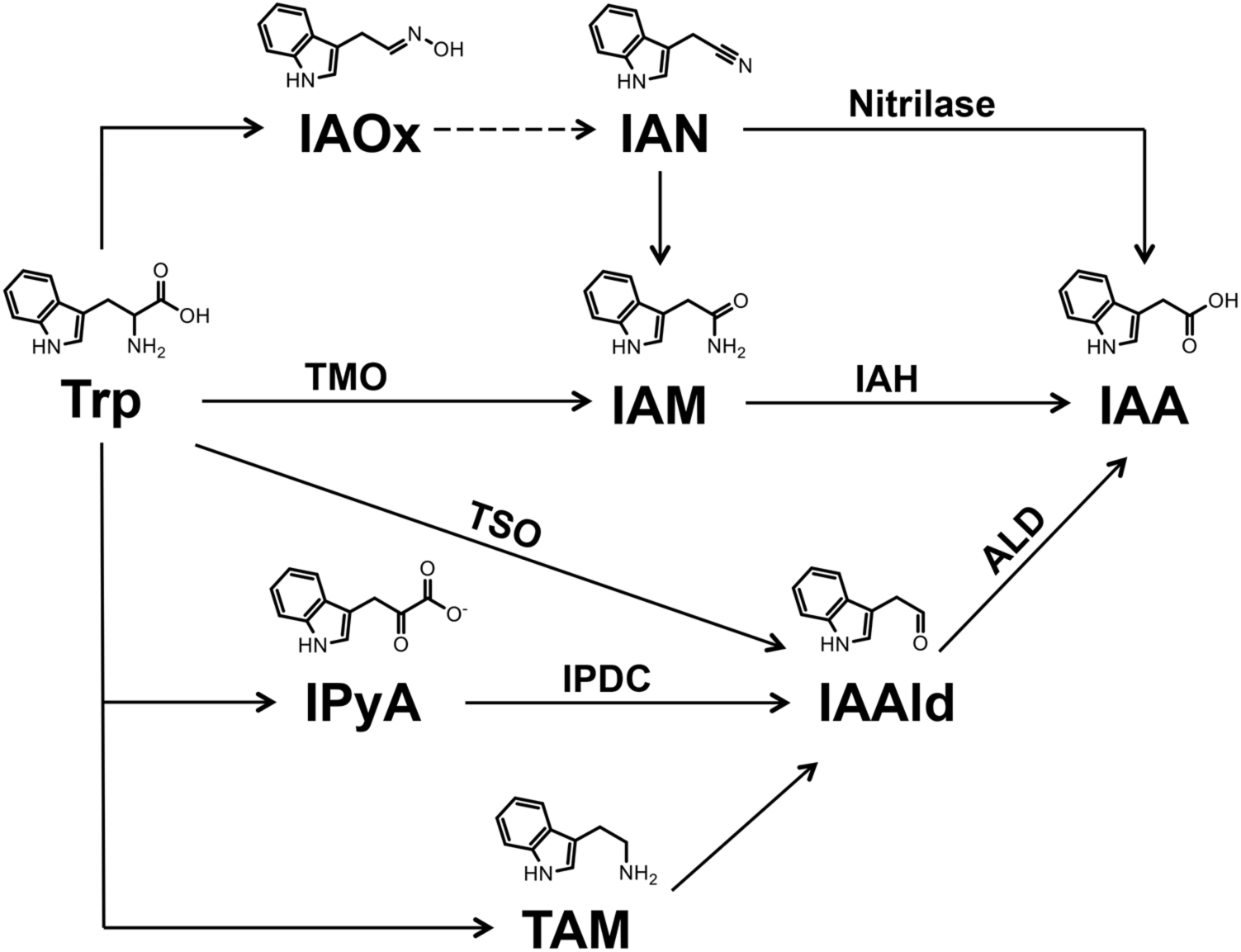
Overview of tryptophan-dependent indole-3 acetic acid (IAA) biosynthesis pathway(s) in bacteria. Enzymes with demonstrated biochemical activities are indicated. Enzyme abbreviations: tryptophan 2-monooxygenase (TMO), indole acetamide hydrolase (IAH), tryptophan side chain oxidase (TSO), indole pyruvate decarboxylase (IPDC) and aldehyde dehydrogenase (ALD). Two ALD enzymes, AldA and AldB, that catalyze conversion of IAAld to IAA are described in this study. Compound abbreviations: tryptophan (Trp), indole-3-acetaldoxime (IAOx), indole-3-acetonitrile (IAN), indole-3-acetamide (IAM), indole-3-pyruvate (IPyA), indole-3-acetaldehyde (IAAld) and tryptamine (TAM).

Auxin is involved in a broad range of growth and developmental processes in plants, including cell division and expansion and responses to a variety of environmental stimuli [20-22]. Auxin is also important in several plant-microbe interactions. For example, IAA produced by plant growth promoting rhizobacteria such as *Azospirillum brasilense* stimulates root growth [23]. IAA also promotes plant cell proliferation during gall formation caused by *Rhizobium radiobacter (*formerly *Agrobacterium tumefaciens)* [24], *Pantoea agglomerans* [18] and *P. savastanoi* [19, 25].

More recently auxin has been shown to promote virulence of *P. syringae* pv. tomato strain DC3000. Exogenous application of auxin enhances disease susceptibility on *Arabidopsis thaliana* [26-28] and transgenic *A. thaliana* lines that over-express the *YUCCA1* auxin biosynthesis gene and accumulate elevated levels of IAA exhibit enhanced susceptibility to DC3000 [29]. Additionally, impairment of auxin signaling in the plant can reduce susceptibility to *P. syringae* pv. tomato and maculicola [27, 30]. Nonetheless, the role of auxin in promoting *P. syringae* virulence remains to be elucidated.

We sought to take advantage of the well-established DC3000-*A. thaliana* interaction to investigate the role of pathogen-derived IAA during pathogenesis. Here, we demonstrate that DC3000 produces IAA and identify an indole-3-acetaldehyde dehydrogenase, AldA, that catalyzes the NAD-dependent formation of IAA from indole-3-acetaldehyde (IAAld). The x-ray crystal structure of AldA provides insight on the biochemical function of this enzyme. We show that disruptions of *aldA* and a close homolog (*aldB*) lead to reduced IAA production in DC3000 and reduced virulence in *A. thaliana*. Furthermore, we explore the mechanism by which pathogen-derived auxin contributes to DC3000 virulence and show that auxin produced by DC3000 suppresses salicylic acid (SA)-mediated defenses in *A. thaliana*.

## Results

### *Pseudomonas syringae* pv. tomato strain DC3000 synthesizes IAA in culture via an indole-3-acetaldehyde intermediate

Many *P. syringae* strains produce IAA in culture, and synthesize elevated levels of IAA when supplemented with Trp [31]; however, it has not been determined whether *P. syringae* pv. tomato strain DC3000 can synthesize IAA. To examine this, we grew DC3000 in Hoitkin-Sinden minimal media containing citrate (HSC) with shaking for 48 hours at 28°C. We chose this media as it is reported to more accurately reflect growth conditions in the intercellular space (e.g. the apoplast) of leaves colonized by *P. syringae* [32]. IAA concentrations in culture supernatants harvested at 24 and 48 hours were determined by LC-MS/MS. As observed for many other *P. syringae* strains, the level of IAA produced by DC3000 was significantly higher (ranging from 100- to 200-fold greater, depending on the experiment) when provided with Trp than in unsupplemented media (Table 1).

**Table 1.** Indole-3-acetic acid (IAA) levels in culture.

| Strain | Supplement<sup>a</sup> | IAA ng/ml<br>24hr<sup>b</sup> | IAA ng/ml<br>48hr<sup>b</sup> |
| --- | --- | --- | --- |
| DC3000 | N/A | 28.9 ± 4.6 | 30.6 ± 3.5 |
| DC3000 | Trp | 2520 ± 245 | 2760 ± 259 |
| DC3000 | IAAld | 3700 ± 189 | 11700 ± 657 |
| DC3000 | IAM | 144 ± 18 | 100 ± 11 |
| DC3000 | IAN | 190<sup>c</sup> ± 7 | 301<sup>c</sup> ± 10 |
| DC3000 | IPyA | 8820<sup>d</sup> ± 331 | 14100<sup>d</sup> ± 285 |
| DC3000 | TAM | 107 ± 20 | 147 ± 10 |

<sup>a</sup>DC3000 cultures grown in Hoitken-Sinden media with 10 mM citrate (HSC) and 0.25 mM of the indicated supplement.
<sup>b</sup>HSC media supplemented with Trp, IAAld, IAM, or TAM accumulated no detectable levels of IAA in the absence of bacteria after 24 or 48 hrs of incubation. Values are average ± SEM (n = 3).
<sup>c</sup>HSC media containing IAN but lacking DC3000 accumulated 111 ng/ml and 124 ng/ml of IAA at 24 and 48 hrs of incubation, respectively.
<sup>d</sup>HSC media supplemented with IPyA but lacking DC3000 accumulated 17,000 ng/ml and 16,300 ng/ml of IAA at 24 hrs and 48 hrs of incubation, respectively. Similar results were obtained in two additional independent experiments.

The observation that DC3000 produces IAA in culture led us to investigate which pathway(s) DC3000 uses to synthesize IAA (Fig 1). The DC3000 genome annotation includes a TMO enzyme (*PSPTO0518; iaaM;* [33]), but the predicted protein exhibits limited amino acid identity to enzymes with demonstrated IAA biosynthetic activity and is more closely related to a second group of TMO homologs that may function in pathways other than IAA synthesis [13]. Thus, it is unclear whether DC3000 uses the IAM pathway to synthesize IAA.

To identify the IAA biosynthetic pathway(s) used by DC3000, we performed IAA precursor feeding experiments using Trp, IAM, IAN, IPyA, TAM, or IAAld and analyzed DC3000 for IAA production by LC-MS/MS. Cultures supplemented with IAM, IAN, and TAM produced small but detectable amounts of IAA compared to cultures grown in HSC alone; however, these levels were relatively low compared to cultures fed with Trp (Table 1). In contrast, at least 100-to 500-fold higher levels of IAA, depending on the incubation time, were produced when DC3000 was grown in media supplemented with IAAld. This indicates that IAAld is an important intermediate for DC3000 IAA synthesis in culture.

The feeding experiments with IPyA were inconclusive, as IPyA is unstable in solution [34] and high amounts of IAA accumulated spontaneously in HSC media containing IPyA, but lacking DC3000 (Table 1). Given the absence of an obvious *ipdc* gene in the DC3000 genome, it is unlikely that DC3000 uses IPyA to synthesize IAAld. Thus, we hypothesize that DC3000 synthesizes IAA via a pathway involving conversion of Trp to IAAld through a TSO activity [35, 36] (Fig 1). We cannot rule out the ability of DC3000 to produce small amounts of IAA through alternative pathways using IAM, IAN and/or TAM; however, based on the results of our feeding studies these pathways do not appear to contribute significantly to IAA synthesis in culture.

### Identification of putative PstDC3000 aldehyde dehydrogenase genes

Our studies indicate that DC3000 synthesizes IAA via one or more pathways that involve IAAld as an intermediate (Fig 1). Thus, we predicted that disrupting the final step, which converts IAAld to IAA, would decrease IAA biosynthesis in DC3000. To investigate this, we sought to identify the gene(s) encoding the enzyme(s) responsible for the conversion of IAAld to IAA. Previously, an *Azospirilum brasilense* mutant (*aldA*) with decreased IAA production was identified and the mutation mapped to a gene encoding a protein with ~80% amino acid identity to an annotated aldehyde dehydrogenase from *Xanthobacter autotrophicus* GJ10 [37]. Aldehyde dehydrogenases (ALDs) generally catalyze the conversion of aldehydes to carboxylic acids [38, 39]. We predicted that a similar enzyme might metabolize IAAld to IAA in DC3000, and thus utilized the amino acid sequences of the ALDs from *A. brasilense* and *X. autotrophicus* to identify putative ALDs in DC3000.

Using BLAST, we identified PSPTO_0728, a putative ALD with ~70% amino acid identity to the ALD from *X. autotrophicus.* We then used the PSPTO_0728 sequence to search the DC3000 genome and identified 5 additional putative ALD homologs, PSPTO_0092, PSPTO_2673, PSPTO_3064, PSPTO_3323, and PSPTO_3644, with ~30-40% amino acid identity to PSPTO_0728. None of these proteins had previously been demonstrated to have dehydrogenase activity, nor were they described as involved in either auxin biosynthesis or DC3000 virulence.

We examined whether these proteins could convert IAAld to IAA by expressing each gene individually in *E. coli*, growing the strains in LB media supplemented with 0.25 mM IAAld, and assaying the resulting strains for IAA production by LC-MS/MS. Background levels of IAA were produced by *E. coli* carrying the empty expression vector (Fig 2), consistent with previous reports [17, 31]. Upon induction of expression of the ALDs from DC3000 (S1 Fig.), we observed increased IAA levels for three of the six proteins. The strains expressing either PSPTO_2673 or PSPTO_3644 showed ~10- and 5-fold increases in IAA levels, respectively (Fig 2A). Cells expressing PSPTO_0092 showed the greatest accumulation of IAA with an ~200-fold increase in IAA over the empty vector control (Fig 2B). Thus, PSPTO_0092, PSPTO_2673, and PSPTO_3644 can convert IAAld to IAA and likely function in DC3000 auxin biosynthesis. We refer to PSPTO_0092, PSPTO_2673, and PSPTO_3644 as AldA, AldB, and AldC, respectively, throughout this study.

**Fig 2.**
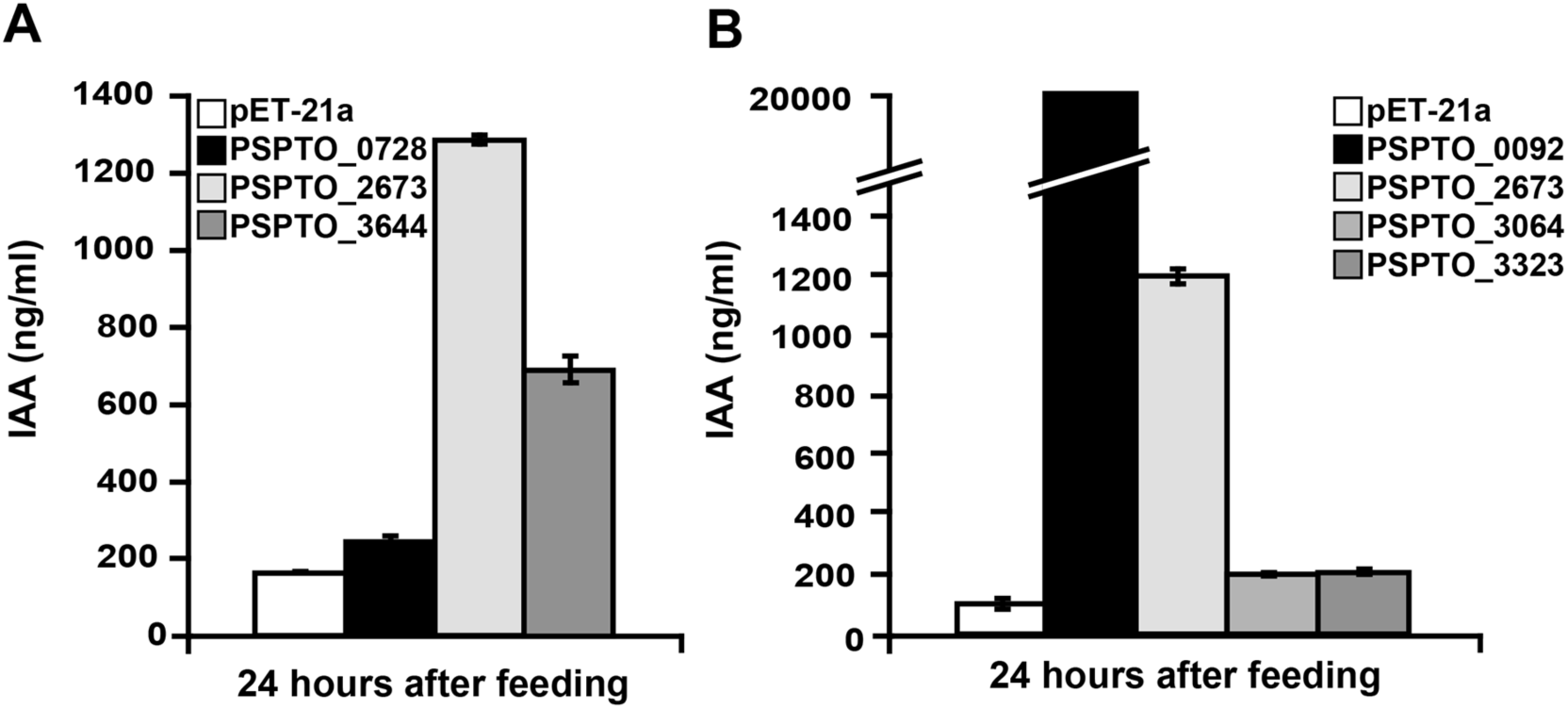
Heterologous expression of putative DC3000 aldehyde dehydrogenases in *E. coli.* DC3000 genes encoding putative aldehyde dehydrogenase proteins were expressed in *E. coli* BL21(DE3) cells. A) Quantification of IAA produced by strains expressing PSPTO_0728, PSPTO_2673, and PSPTO_3644 and pET-21a as a negative control. B) Quantification of IAA produced by strains expressing PSPTO_0092, PSPTO_3064, and PSPTO_3323. PSPTO_2673 was included as a control for comparison to panel A. IAA levels were measured in supernatants 24 hrs post-induction with addition of 0.25 mM IAAld. Values are an average ± SEM (n=3). Similar results were obtained from two additional independent experiments.

### Biochemical analysis of putative IAAld dehydrogenas

Based on sequence comparisons, AldA-C are members of the aldehyde dehydrogenase enzyme superfamily [38, 39] (S2 Fig.). To examine the biochemical activity of the three putative ALDs from DC3000, these proteins were expressed in *E. coli* as a N-terminal hexahistidine-tagged proteins and purified by nickel-affinity and size-exclusion chromatographies. Each of the putative ALDs was isolated with a monomer M_r_~56 kDa, as determined by SDS-PAGE (S3A Fig.), which corresponds to the estimated molecular weights of AldA (M_r_ = 52.7 kDa), AldB (M_r_ = 53.1 kDa) and AldC (M_r_ = 51.8 kDa) plus the addition of a His-tag. Size-exclusion chromatography of AldA and AldB indicates that each protein functions as a tetramer and that AldC is dimeric (S3B Fig.).

In vitro assays of purified AldA, AldB and AldC using IAAld with either NAD^+^ or NADP^+^ as substrates confirm the major activity of AldA as that of an IAAld dehydrogenase, as each protein converted NAD(P)^+^ to NAD(P)H only in the presence of the IAAld (S3B Fig.). Each Ald used NAD+ with a 10- to 40-fold preference versus NADP^+^, but AldA had a specific activity (3.52 μmol min^-1^ mg protein^-1^) using IAAld as a substrate that was 100- and 800-fold higher than AldB and AldC, respectively. AldA-C displayed no changes in specific activities in the presence of calcium, magnesium, manganese, cobalt, nickel, and copper, which suggests that these proteins function as non-metallo NAD^+^-dependent ALDs. None of the three Alds showed detectable activity with IAA (at 1 mM) and NADH (at 200 μM), indicating a clear preference for the formation of IAA compared to the reverse reaction. Kinetic analysis showed that AldA had a catalytic efficiency (*k*_cat_/*K*_m_) with IAAld as a substrate that was 130- and 710-fold higher than AldB and AldC, respectively (S1 Table). AldA also showed more than a 300-fold higher *k*_cat_/*K*_m_ with NAD^+^ compared to NADP^+^. A similar cofactor preference was observed for AldB and AldC. The low activities of AldB and AldC did not allow for accurate determination of kinetic parameters for NADP^+^. These biochemical comparisons suggest that AldA functions as an IAAld dehydrogenase and that AldB and AldC likely prefer other aldehyde substrates in vivo.

### Overall three-dimensional structure of AldA

To explore the molecular basis of IAAld dehydrogenase activity of AldA, its three-dimensional structure was determined by X-ray crystallography. The X-ray crystal structures of AldA in the apoenzyme, NAD^+^ complex, and NAD^+^•IAA complex forms were determined (S2 Table). In each structure, two AldA monomers were in the asymmetric unit and packed to form a dimer, which then form a tetramer by crystallographic symmetry (Fig 3A). The interface between two monomers buries ~2,450 Å^2^ of surface area with a ~3,800 Å^2^ interface between each of the dimer units. The overall fold of AldA shares structural similarity with ALDH2-3 (4PXL; 1.2 Å r.m.s.d. for ~480 C_α_-atoms; 46% identity) and ALDH2-6 (4PZ2; 1.3 Å r.m.s.d. for ~484 C_α_-atoms; 46% identity) from *Zea mays*, along with multiple human ALD structures (1.4 - 1.5 Å r.m.s.d. for ~400 C_α_-atoms; 43-46% identity) [40, 41]. The AldA monomer adopts a canonical aldehyde dehydrogenase fold (Fig 3B), which contains an NAD^+^-binding domain with a Rossmann-fold motif of a central β-sheet (β10-β9-β8-β11-β12) surrounded by α-helices, a mixed α/β domain with the catalytic cysteine residue (Cys302), and an oligomerization domain with a protruding β-sheet (β6-β7-β23).

**Fig 3.**
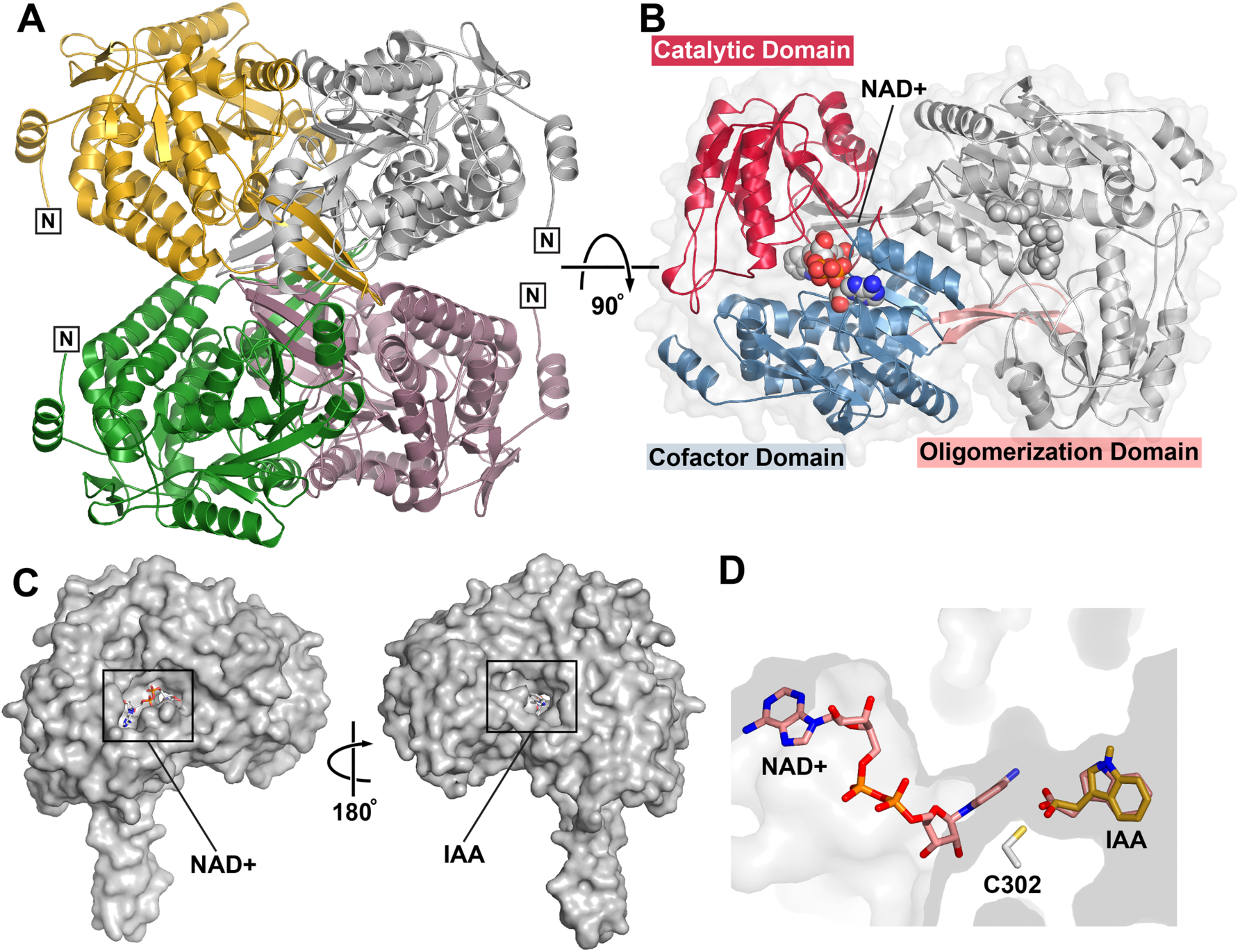
Overall structure of AldA. A) The AldA tetramer is shown as a ribbon tracing with each subunit differentially colored. Two subunits (gold and white) were in the asymmetric unit of the crystal with the other two subunits (green and rose) related by crystallographic symmetry. N-termini are labeled. B) Domain organization of the AldA monomer. The view is rotated 90° relative to panel A and shows the two subunits in the asymmetric unit. The catalytic (red), cofactor binding (blue), and oligomerization (rose) domains are highlighted in one monomer. The position of NAD^+^ (space-filling model) is indicated. C) Substrate binding sites on opposite sides of the AldA monomer. The two views of an AldA monomer are rotated 180° and show the locations of the NAD(H) and IAAld/IAA binding sites on each face of the monomer. D) Ligand binding tunnel. The positions of NAD^+^ (rose) and IAA (gold) in the tunnel (grey surface) relative to the catalytic cysteine (Cys302) are shown. The position of docked IAAld (rose), which overlaps with IAA, is indicated.

The AldA•NAD^+^ and AldA•NAD^+^•IAA crystal structures define the position of the active site between the catalytic and cofactor binding domains (Fig 3B-C). Although the ligand binding sites occupy two separate pockets on opposite sides of the monomer (Fig 3C), both sites are linked by a ~25 Å tunnel that places the reactive groups of the co-substrates in proximity to Cys302 (Fig 3D). Comparison of the AldA crystal structures suggests that ligand binding results in structural changes that order the active site (S3D Fig.). The α11-β14 loop (residues 297-305), which contains Cys302, is disordered in the apoenzyme structure and has average temperature factors ~1.8-fold higher than surrounding residues. Likewise, a ~50 amino acid region of the catalytic domain (residues 348-397; α13-β15-β16-α15-β17-β19) is disordered in the apoenzyme structure and displays elevated B-factors in ligand bound structures.

### Structure of the AldA active site

Unambiguous electron density in the AldA•NAD^+^ and AldA•NAD^+^•IAA crystal structures identifies the respective ligand binding sites (Fig 4A). In the NAD^+^ binding site, the cofactor is bound in a hydrophobic tunnel (Fig 4B). The adenine ring of NAD^+^ lies in an apolar region that provides multiple van der Waals contacts. The adenine ring also forms two hydrogen bonds between the hydroxyl group of Tyr255 and a water. The adenine-ribose rings provide extensive polar interactions with AldA. The 2’-hydroxyl hydrogen bonds with Lys191 and Glu194. Interactions with Ser193, Ser245, and Trp167 position the phosphate backbone in the binding site. The nicotinamide-ribose forms a bidentate interaction with Glu401 and the nicotinamide ring is bound by a water-mediated interaction with Thr243 and through a hydrogen bond from Glu267. Sequence comparisons show a conserved NAD+ binding site in AldA, AldB and AldC (S2 Fig.). These interactions place the nicotinamide ring in proximity to the invariant catalytic cysteine (Cys302 in AldA) [38].

**Fig 4.**
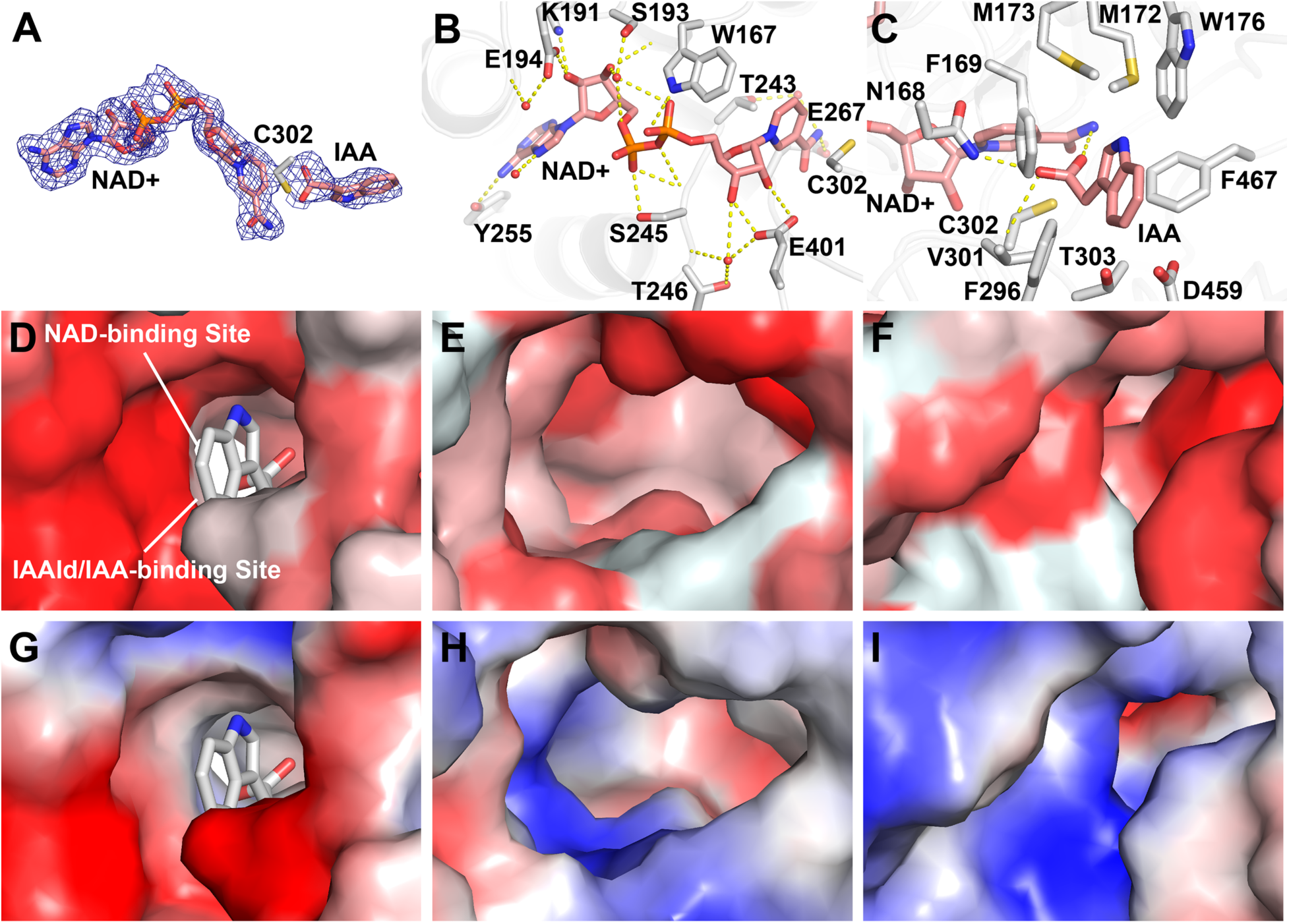
Substrate and cofactor binding sites of AldA. A) Electron density of NAD^+^ and IAA. The 2F_o_-F_c_ omit map (1.5 s) for NAD^+^ and IAA is shown. B) NAD(H) binding site. Side-chains of residues interacting with NAD^+^ (rose) are shown as stick-renderings. Waters interacting with the cofactor are shown as red spheres. Hydrogen bonds are indicated by dotted lines. C) IAAld/IAA binding site. NAD^+^, IAA, and side-chains are shown as stick-renderings with dotted lines indicating hydrogen bonds. D-F) Hydrophobicity of the substrate binding sites of AldA (panel D), AldB (panel E), and AldC (panel F). Homology models of AldB and AldC were generated based on the x-ray structure of AldA. Hydrophobicity was calculated using the Color-h script in PyMol. Darkest red indicates strongest hydrophobicity to white as the most polar. G-I) Electrostatic surface of the substrate binding sites of AldA (panel G), AldB (panel H), and AldC (panel I). Electrostatic surface charge was generated using the APBS plugin in PyMol with red = acidic and blue = basic.

Crystallization of a ‘dead-end’ complex (i.e., AldA•NAD^+^•IAA) provides insight on the IAAld binding site (Fig 4A and C). Electron density was observed near the reactive Cys302 and modeled as IAA for refinement. In contrast to NAD(H) binding, the IAAld/IAA site is formed predominantly by apolar residues. The carboxylic acid of IAA forms hydrogen bonds with the sulfhydryl group of Cys302, the amide side-chain of Asn168, and the backbone nitrogen of Cys302. Multiple aromatic and apolar residues, including Phe169, Met173, Trp176, Val301, and Phe467, surround the indole moiety. Computational docking of IAAld into the active site yielded a solution that matched the crystallographically observed position of IAA (Fig 3D). The docked IAAld overlays with IAA and positions the reactive aldehyde group of the substrate near Cys302 for subsequent catalysis.

To understand the different activity with IAAld displayed by the three ALDs, homology models of AldB and AldC based on the AldA structure were generated. Although the NAD(H) binding sites of AldA-C are highly conserved, the residues in the aldehyde binding site of each enzyme displays greater variability (S2 Fig). Compared to AldA, sequence differences in AldB and AldC alter the hydrophobicity, electrostatics, and surface shape of the site (Fig 4D-I). For example, the calculated hydrophobicity values of the IAAld/IAA binding site are 7.51 in AldA, - 2.99 in AldB, and 2.78 in AldC (Fig 4D-F). Likewise, the surface electrostatics of AldB and AldC are more basic than AldA (Fig 4G-I). In addition, the shape of the site in each enzyme differs. The largely apolar IAAld/IAA binding site of AldA best fits the substrate molecule. Amino acid changes in the AldB may widen the substrate binding pocket. The wider and more basic nature of this site likely reduces catalytic efficiency of AldB with IAAld. Whereas, substitutions in the AldC substrate binding site likely constrict access to the catalytic cysteine and result in the even lower activity of this enzyme with IAAld. Thus, structural differences in the substrate binding sites of these ALD result in the preference of AldA for IAAld.

### IAA production is disrupted in DC3000 *ald* mutants

To study the role of these ALDs in DC3000 IAA biosynthesis, we generated plasmid disruption mutants in *aldA* (*PSPTO_0092)*, *aldB* (*PSPTO_2673)* and *aldC* (*PSPTO_3644)* (Fig S4). The mutant strains were not notably different from DC3000, other than exhibiting a small but significant reduction in growth in NYG or HSC media (S4 Fig. E and F). We monitored the ability of each mutant strain to produce IAA in culture when provided with IAAld. Only two mutants displayed reduced levels of IAA when compared to DC3000 (Fig 5A). The *aldA* mutant displayed a ~75% reduction in IAA levels compared with DC3000, whereas the *aldB* mutants exhibited a ~15% reduction in IAA levels. These results indicate that AldA and AldB proteins contribute to IAA synthesis in DC3000, but that AldC does not.

**Fig 5.**
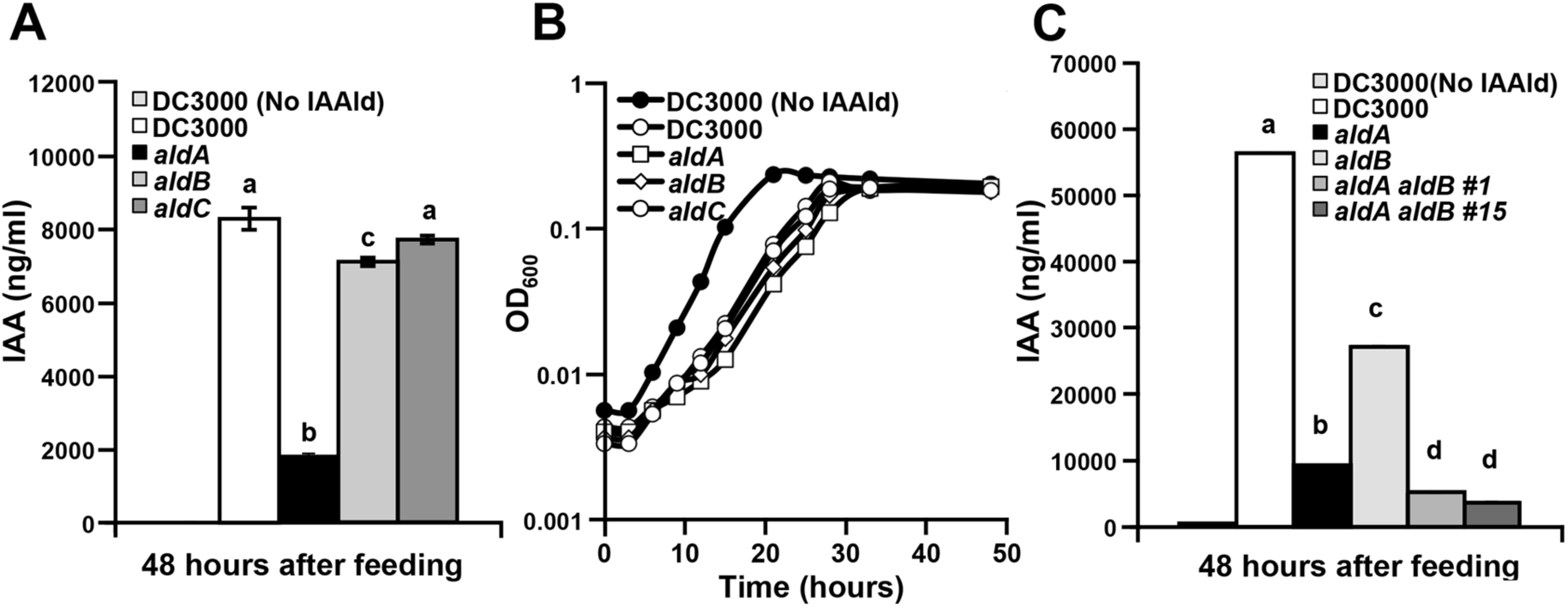
Quantification of IAA production in DC3000 *ald* mutants. A) Measurement of IAA accumulation in supernatants of DC3000 *ald* single mutants grown for 48 hrs in HSC media supplemented with 0.25 mM IAAld. B) Growth of *ald* mutants in HSC media supplemented with 0.25 mM IAAld. Cultures were used to quantify IAA shown in panel A. C) Measurement of IAA accumulation in supernatants of two independent *aldA aldB* double mutants grown for 48 hrs in HSC media supplemented with 0.25 mM IAAld. For panels A-C, values are an average of three biological replicates ± SEM (error bars too small to see in panels B and C). Letters indicate significant difference between samples within a given time point (*p*<0.05).

Interestingly, DC3000 exhibited reduced growth rates in HSC media supplemented with IAAld compared to DC3000 grown in HSC alone (Fig 5B). This could be due to a toxic effect of IAAld at the given concentration (0.25 mM). All three *ald* mutant strains also displayed a similar reduction in growth rates in HSC media supplemented with IAAld.

### DC3000 IAA biosynthesis mutants exhibit reduced virulence on *Arabidopsis thaliana*

Previous studies indicate that auxin promotes susceptibility to DC3000 and *P. syringae* pv. maculicola ES4326 [26-30]; however, it is unknown whether auxin produced by these strains contributes to their virulence. To examine this, we assayed the *aldA* and *aldB* mutants for altered virulence on *A. thaliana* plants. DC3000 grew to high levels when infiltrated into *A. thaliana* plants (Fig 6A), while the *aldA* and *aldB* mutants exhibited a ~5-fold reduction in growth. Surface inoculation experiments were also performed to monitor development of disease symptoms. Plants dip-inoculated with DC3000 exhibited characteristic disease symptoms consisting of many individual water-soaked lesions surrounded by yellowing of the leaf (chlorosis) (Fig 6B-C). Plants infected with the *aldA* mutant displayed reduced disease symptom severity compared to DC3000, manifested primarily as a decrease in the percentage of leaves developing high levels of chlorosis. Both the reduced IAA synthesis and reduced virulence phenotypes of the *aldA* mutant were complemented by introduction of the wild-type *aldA* genomic clone (S5 Fig.), indicating that DC3000-derived IAA contributes to virulence. Plants infected with the *aldB* mutant also displayed a reduction in symptom severity, although to a lesser degree than plants infected with the *aldA* mutant (Fig 6B-C).

**Fig 6.**
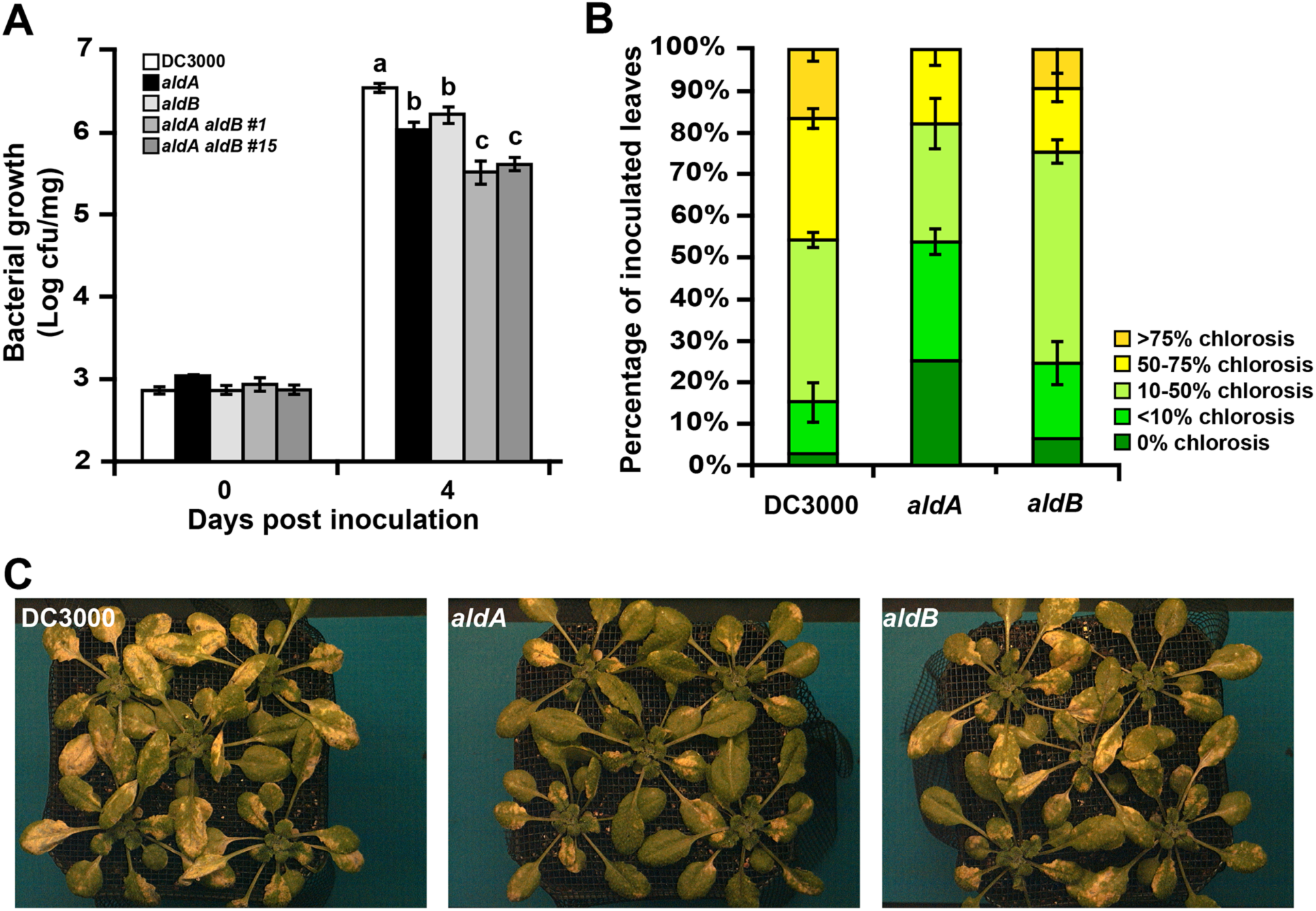
Growth and symptom production of *ald* mutants on. *A. thaliana.* A) Growth of DC3000 and *ald* mutants following syringe infiltration of *A. thaliana* (OD_600_ =1×10^−4^). Similar results were seen in two additional experiments. Letters indicate significant difference between samples within a given time point (*p*<0.05). B) Disease symptom severity 4 days after dip inoculation with *ald* mutants. Disease symptom severity was quantified as the average percentage of affected leaves per plant exhibiting the indicated amount of chlorosis. 10 plants were assayed for each treatment. Results are plotted as the average percentage of leaves from each genotype exhibiting the indicated degree of chlorosis. C) Photographs taken 4 days after dip inoculation. Plants shown were used to quantify disease symptom severity in panel B. Similar results were obtained in two additional experiments.

We tested whether the *ald* genes have an additive effect on IAA synthesis and virulence by generating an *aldA aldB* double mutant in DC3000. We monitored the ability of two independent double mutant strains to produce IAA in culture when fed with IAAld, and observed that IAA production was significantly lower in *aldA aldB* double mutants than in either single mutant (Fig 5C). The *aldA aldB* double mutant also exhibited a further reduction in bacterial growth on *A. thaliana* plants compared to the single mutants (Fig 6A). The additive nature of these mutant phenotypes suggests that AldA and AldB contribute to DC3000 IAA biosynthesis and virulence in a partially redundant manner. As the *aldA aldB* double mutant exhibits reduced grown in minimal media (S4 Fig. E&F), the additive effect on growth in planta may reflect a more general role for ALD activity in *P. syringae* metabolism.

### Pathogen-derived IAA suppresses SA-mediated defenses

IAA may contribute to pathogenesis by suppressing host defenses mediated by the defense hormone SA [27, 42]. We hypothesized that if pathogen-derived IAA promotes pathogen growth in planta by suppressing SA-mediated defenses, then the reduced growth of the DC3000 *ald* mutants in planta would be associated with elevated SA-mediated defenses due to an impairment in the ability to suppress SA-mediated defenses. To investigate this, we monitored the expression of *PR1*, a commonly used marker for SA-mediated defenses in *A. thaliana* [29], in plants infected with wild-type DC3000 and the *aldA* and *aldB* mutants 24 hours after inoculation. *PR1* expression was induced by 24 hrs in plants infected DC3000 compared to mock treatment (Fig 7A). Expression of *PR1* was significantly higher in plants infected with the *aldA* mutant. There was also a significant increase in *PR1* expression in plants infected with the *aldB* mutant; however, this increase was not as large as observed for the *aldA* mutant. These results suggest that DC3000-derived IAA is required for normal virulence via a mechanisms involving suppression of SA-mediated defenses.

**Fig 7.**
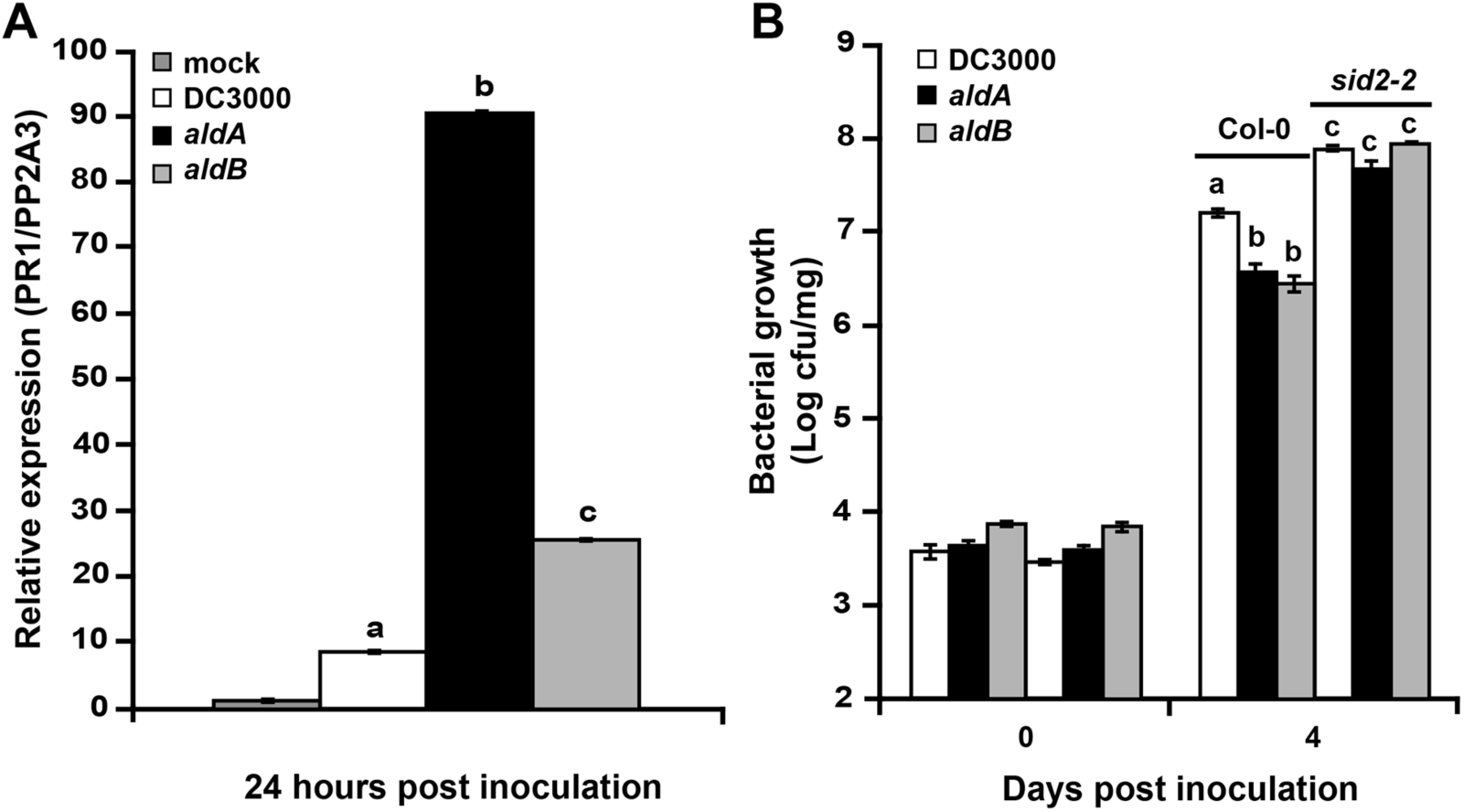
*PR1* expression in plants inoculated by *ald* muants and growth of *ald* mutants on SA - deficient *sid2*-*2* plants. A) *PR1* expression in Col-0 plants at 24 hrs following syringe infiltration (OD_600_ = 1×10^−5^). Significant elevation of *PR1* expression in *aldA*-infected plants was observed in three independent experiments, and in two experiments for *aldB*-infected plants. B) Growth of *ald* mutants on wild type *A. thaliana* (Col-0) and *sid2*-*2* mutant plants following syringe infiltration (OD_600nm_ = 1×10^−4^). Similar growth differences were observed in two additional experiments. Letters indicate significant difference between samples within a given time point (*p*<0.05).

Given these findings, we predicted that the growth of the *ald* mutants would be restored to wild-type levels on *A. thaliana* mutants with impaired SA-mediated defenses. To test this, we inoculated the *sid2*-*2* mutant, which carries a mutation in the *ICS1* SA biosynthesis gene [43], with DC3000 and the *ald* mutants and monitored bacterial growth. Wild-type DC3000 grew to higher levels in *sid2*-*2* mutants plants than in wild-type Col-0 (Fig 7B), consistent with previous reports that the *sid2*-*2* mutant exhibits increased disease susceptibility to *P. syringae* [29, 43].

Consistent with our earlier results, the *aldA* and *aldB* mutants exhibited significantly reduced growth on wild-type plants compared to DC3000; however, each mutant grew to levels comparable to wild-type DC3000 on *sid2*-*2* plants (Fig 7B). Thus, reduced growth of the *ald* mutants is restored to normal levels in plants impaired for SA-mediated defenses. These results suggest that DC3000-derived IAA promotes pathogen virulence by suppressing SA-mediated defenses.

## Discussion

Natural (i.e., IAA) and synthetic (i.e., naphthaleneacetic acid and the herbicide 2,4-dichlorophenoxyacetic acid) auxins can promote virulence of *P. syringae* [26, 27, 29, 30]. Although many plant-associated bacteria can synthesize IAA in culture [11, 12], the role of pathogen-produced IAA in interactions between non-gall-inducing *P. syringae* strains and their hosts is not clear. We investigated this by examining the role of IAA synthesis by *P. syringae* strain DC3000 during pathogenesis of *A. thaliana.* In this work, we demonstrate that DC3000 synthesizes IAA in culture when fed with the either Trp or IAAld and identify an indole acetylaldehyde dehydrogenase, AldA, that converts IAAld to IAA. Based on our biochemical and genetic analyses, AldA is responsible for the majority of IAA synthesis in culture and is required for full virulence of DC3000 on *A. thaliana* plants. These results suggest that AldA-dependent synthesis of IAA plays an important role during pathogenesis.

### DC3000 synthesizes IAA in culture via an IAAld intermediate

Using precursor feeding studies we determined that DC3000 synthesizes IAA in culture via a biosynthetic pathway that utilizes IAAld as an intermediate. To further investigate IAA synthesis, a reverse genetic approach identified a family of ALDs that catalyze the reduction of IAAld to IAA. Of this family, AldA is the enzyme primarily responsible for IAA synthesis from IAAld in culture (Fig 5). A second enzyme, AldB also contributes to IAA synthesis, but seems less important than AldA, based both on its lower activity in vitro (S3C Fig.) and on the observation that IAA production by the *aldB* mutant is only moderately reduced (Fig 5). The two enzymes appear to function redundantly in culture, as IAA synthesis is further reduced in the *aldA aldB* double mutant; however, the double mutant still accumulates some IAA in cultures fed with IAAld, which indicates there may be one or more additional genes encoding IAAld dehydrogenase activity.

### AldA is an Indole Acetylaldehyde Dehydrogenase

Biochemically, aldehyde dehydrogenases (ALDs) are a large enzyme superfamily that convert aldehydes to carboxylic acids on a broad array of molecules [38, 39, 44]. In diverse organisms, multiple ALDs function in various metabolic pathways and provide house-keeping functions, such as the detoxification of reactive aldehydes produced by lipid peroxidation. As with other enzyme superfamilies, the aldehyde dehydrogenases are an excellent example of how evolution of different substrate specificity while retaining common reaction chemistry leads to functional diversity and tailoring of biological function [45]. This appears to be the case for the ALDs in DC3000, as AldA has a specialized role in IAA biosynthesis and pathogenesis that is distinct from AldB and AldC.

Structurally, AldA shares the same overall three-dimensional fold as other ALDs (Fig 3) and functions as an NAD(H)-dependent enzyme (S3C Fig.; S1 Table). Although AldA shares ~40% amino acid identity with both AldB and AldC (S2 Fig.), kinetic analysis of AldA demonstrates a distinct preference for IAAld as a substrate compared to the other two enzymes. The x-ray crystal structure of AldA in complex with NAD^+^ and IAA reveals the molecular basis for the activity of this protein (Fig 4). In the reaction sequence catalyzed by AldA, substrate binding leads to conformational changes that order the active site for catalysis (S3D Fig.). The chemical mechanism would proceed as described for other aldehyde dehydrogenases [46]. For conversion of IAAld to IAA, the active site cysteine (Cys302) acts as a nucleophile to attack the substrate aldehyde moiety. This leads to formation of a covalent intermediate. Subsequence transfer of a hydride from the substrate to NAD^+^ and nucleophilc attack by an activated water molecule on the resulting carbonyl of the intermediate releases the carboxylic acid product with the thiol acting as a leaving group.

Comparison of the structure and sequence of AldA with AldB and AldC shows how changes alter the size, shape, hydrophobicity, and electrostatics of the binding pocket (Fig 4D-I). Thus, the evolution of the AldA substrate binding site leads to a preference for IAAld. Additional studies are needed to identify the preferred substrates of AldB and AldC. Overall, the biochemical and structural data presented here indicate that in *P. syringae* strain DC3000 AldA functions as an IAAld dehydrogenase in IAA biosynthesis. This is the first identified in either plants or microbes and suggests that the evolution of different metabolic routes to IAA synthesis can be exploited by microbial plant pathogens.

### The DC3000 IAA biosynthesis pathway

We propose that AldA-dependent IAA synthesis in DC3000 involves the direct conversion of Trp to IAAld through TSO activity (Fig 1), as the DC3000 genome does not encode an obvious IPDC, nor do our feeding studies implicate TAM as an intermediate (Table 1). The TSO pathway, which has been reported in several *P. fluorescens* strains [11], is not well characterized. A Tn5 mutant lacking TSO activity was identified in *P. fluorescens* strain CHA0 [35]; however, a gene encoding this activity has not been described. Future investigation of TSO activity in DC3000 will provide additional insight into IAA synthesis in *P. syringae* and other bacteria.

We also investigated the hypothesis that DC3000 utilizes the IAM pathway, as this pathway is used by other IAA producing bacteria, including several *Pseudomonas* strains [12, 31]. Neither our feeding studies, nor recent bioinformatic and genetic analyses provide support for the existence of an IAM pathway in DC3000. Patten et al. [13] noted that *PSPTO_0518*, which is annotated as encoding a TMO (Fig 1; *iaaM*, [33]; http://www.pseudomonas.com), shares only ~30% amino acid identity with enzymes with demonstrated TMO activity. PSPTO_0518 is more closely related to a second family of monooxygenases that may function in other pathways [13]. Further, our observation that mutation of *PSPTO_0518* does not alter accumulation of IAA in cultures fed with Trp provides additional evidence for the absence of the IAM pathway in DC3000 (A. Mutka, E. Mellgren and B. Kunkel, unpublished). Likewise, our feeding studies do not implicate the IAN pathway as a major contributor to IAA synthesis in DC3000 (Table 1).

Many Pseudomonads, including *P. syringae*, *P. fluorescens*, *P. putida*, and *P. aeruginosa*, have genes predicted to encode proteins with ~90-95% sequence identity to AldA, including a nearly invariant conserved IAAld binding site. A survey of The Pseudomonas Genome Data Base (www.pseudomonas.com) revealed that AldA homologs are much more common in these genomes than TMO, which is only found in a few *P. syringae* or *P. savastanoi* strains [13]. Thus, we speculate that the AldA-dependent IAA biosynthesis pathway is the predominant IAA synthesis pathway in Pseudomonads. The role of IAA production in the biology of these microbes is yet to be elucidated; however, in the case of plant-associated bacteria, modification of the biology of their plant hosts seems likely. Alternatively, or additionally, IAA may be involved in signaling with other microbes in the soil or leaf epiphytic community [12, 19].

### What is the role of Ald(A)-dependent IAA synthesis in planta?

Our observation that the *aldA*, *aldB* and the *aldA aldB* double mutant strains exhibited both reduced growth and reduced disease symptom production on *A. thaliana* plants (Fig 6) suggests that AldA and AldB play important roles during pathogenesis. Although the single mutants exhibited slightly reduced growth in culture (S4 Fig.), the fact they grow to high levels in *sid2* plants (Fig 7) indicates that the reduced growth of these strains in wild type plants does not reflect a general growth defect. Thus, both Ald activities contribute to DC3000 virulence on *A. thaliana.* Kinetic comparisons indicate that AldA is more specific than AldB for IAAld; however, differences in protein expression in the microbe (i.e., high levels of AldB) could allow for the less efficient enzyme to contribute to IAAld conversion to IAA. We have not demonstrated that one or both enzymes catalyze IAA production in planta, as it is technically difficult to distinguish pathogen-derived from plant-derived auxin in plant tissue. However, it is reasonable to expect that this is the case, as both Trp and IAAld are present in significant amounts in *A. thaliana* tissue [47, 48].

It is interesting to note that, even though both *ald* mutants exhibit a similar (~5-fold) reduction in growth in planta (Fig 6A, 7B), the reduction in disease symptom severity caused by *aldA* was much more pronounced than for *aldB*, suggesting that AldA-dependent IAA synthesis is important in promoting disease symptom development. Given that our biochemical studies suggest that AldB may not use IAAld as a substrate (S3C Fig.), the exact role of AldB during infection is presently not clear. It is possible that reduction of some other aldehyde by AldB in planta contributes to virulence.

Our observation that plants infected with the *aldA* mutant express elevated levels of *PR1* mRNA (Fig 7A) suggests that pathogen-derived IAA promotes virulence by suppressing SA-mediated defenses. Consistent with this, we also showed that growth of the *aldA* mutant is restored to wild-type levels on SA-deficient plants (Fig 7B). These findings agree with results from earlier studies demonstrating that exogenous application of auxin down-regulated SA-mediated defenses [27, 49].

### IAA plays multiple roles during pathogenesis

The finding that pathogen-derived IAA promotes DC3000 virulence by suppressing SA-mediated defenses contrasts with results from our previous studies with transgenic plants that overexpress the *YUCCA1 (YUC1)* IAA biosynthesis gene and accumulate elevated levels of IAA [50]. We observed that *YUC1* overexpressing plants exhibited increased susceptibility to DC3000, but that neither SA accumulation nor SA-responsive gene expression was suppressed in these plants [29]. Further, plants carrying both the *YUC1* overexpression construct and the *sid2* mutation exhibited additive effects of enhanced susceptibility due to both elevated IAA and impaired SA-mediated defenses. These results suggest that in these plants, IAA promotes pathogen growth through mechanism that functions independently of suppression of SA-mediated defenses [29]. The apparent discrepancy between these studies can be resolved by proposing that: 1) auxin promotes DC3000 virulence via multiple different mechanisms, and 2) pathogen-derived auxin and plant-derived auxin play different roles during pathogenesis (Fig 8). Our data suggest that the stimulatory effect of AldA-dependent DC3000-synthesized IAA on virulence acts via suppressing SA-mediated defense signaling, while auxin produced by the plant (e.g. *YUC1*-dependent) promotes pathogen growth via a mechanism that acts independently or down-stream of SA-mediated defenses. Future studies examining the impact of the source, and possibly also the form, of auxin during pathogenesis will provide important insight into the roles of auxin in promoting disease development by DC3000. It will also be of interest to investigate whether auxin plays multiple roles in other plant-microbe interactions.

**Figure 8.**
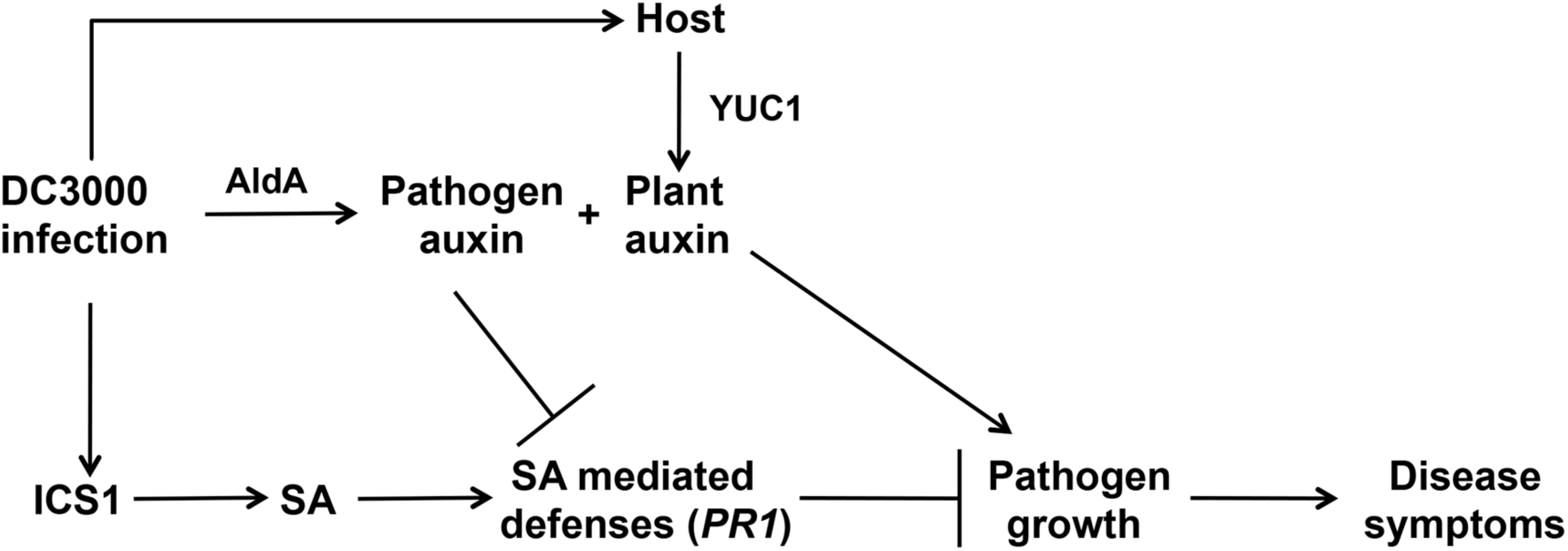
IAA promotes pathogenesis via multiple mechanisms. DC3000 synthesizes IAA via the activity of the aldehyde dehydrogenase AldA. The DC3000 *aldA* mutant exhibits reduced virulence on *A. thaliana* and plants infected with *aldA* express elevated SA-mediated defenses, suggesting that pathogen-derived IAA promotes virulence by suppressing SA-mediated defenses. Previous studies have shown that exogenous application of auxin promotes disease [26, 30] and inhibits SA-mediated defenses [27], but that in transgenic plants overexpressing the *YUCCA1* (*YUC1*) IAA biosynthesis gene and that accumulate elevated IAA, increased susceptibility to DC3000 occurs via a mechanism that does not involve suppression of SA-mediated defenses [29]. Together, these observations suggest that pathogen-produced auxin and plant-produced auxin promote disease via different mechanisms. SA, salicylic acid; *ICS1/SID2*, *ISOCHORISMATE SYNTHASE 1*; *PR1*, *PATHOGENESIS RELATED 1*

## Materials and Methods

### Bacterial strains and plasmids

The bacterial strains and plasmids used in this study are summarized in Supplemental Table S3. *P. syringae* strain DC3000 wild-type and mutant strains were grown on Nutrient Yeast Glycerol Medium (NYG) [51] or Hoitkin Sinden (HS) Medium (with appropriate carbon sources added) at 28°C. HS was prepared as described in [52]. *Escherichia coli* strains were maintained on Luria Broth (LB) medium at 37°C. Antibiotics used for selection of *P. syringae* strains include: rifampicin (Rif, 100 μg mL^-1^), kanamycin (Kan, 25 μg mL^-1^), and tetracycline (Tet, 16 μg mL^-1^). Antibiotics used for selection of *E. coli* strains were ampicillin (Amp, 100 μg mL^-1^), Kan (25 μg mL^-1^) and chloramphenicol (Cm, 20 μg mL^-1^).

A modified version of the pJP5603 suicide vector [53], pJP5603-Tet, in which the Kan^R^ cassette was replaced with the Tet^R^ gene, was constructed for generation of double insertion/disruption mutants. The pJP5603-Tet vector was made by digesting pJP5603 with XbaI and BglII to release the ~1.3kb Kan^R^ cassette, and an ~2.9kb XbaI and BglII fragment containing the Tet^R^ gene from pME6031 was inserted in its place.

### Quantification of indole-3-acetic acid (IAA) production in culture

*P. syringae* strains were grown in NYG medium with Rif in overnight cultures. Cells were collected by centrifugation from each overnight culture, washed twice with 10 mM MgCl_2_, re-suspended at a density of ~1 × 10^7^ cells mL^-1^ in HS minimal media containing 10 mM citrate and incubated with shaking for 48 hrs at 28 °C. The culture medium was supplemented with 0.25 mM Trp, IAM, IAN, TAM, or IAAld, as indicated. One mL samples were taken at 24 and 48 hrs after incubation, centrifuged to pellet the cells and the resulting supernatants frozen in liquid nitrogen and stored at -80 °C. Growth of cultures was monitored by reading the OD_600_ at regular intervals with a spectrophotometer. The samples were analyzed for IAA production by LC-MS/MS [54].

### Bioinformatics, nucleotide sequences, and accession numbers

BLASTP searches were performed using the National Center for Biotechnology Information (NCBI) server to search non-redundant databases for *P. syringae* DC3000-specific sequences. *P. syringae* strain DC3000 sequence information was obtained from Kyoto Encyclopedia of Genes and Genomes (KEGG; www.genome.jp/kegg) and the Pseudomonas-Plant Interaction website (PPI; www.pseudomonas-syringae.org). Accession numbers for genes used in this study are: aldehyde dehydrogenase (AldA) from *A. brasilense:* AY850388; chloroacetaldehyde dehydrogenase (AldA) from *X. autotrophicus*: AF029733; DC3000 PSPTO_0092 (AldA): NP_789951.1; DC3000 PSPTO_2673 (AldB): NP_792480.1; DC3000 PSPTO_3644 (AldC): NP_793419.1.

### Expression of *P. syringae* putative aldehyde dehydrogenase genes in *E. coli*

To make the pET21a-0092 (AldA) expression plasmid, the full-length coding sequence (CDS) from *PSPTO_0092* was amplified from DC3000 genomic DNA with primers 0092NdeI F and 0092XhoIR (S4 Table). The resulting ~ 1.5 kb PCR fragment was cloned into the pBlunt II-TOPO vector (Invitrogen), transformed into *E.coli* DH5α and plated on LB media containing Kan. The resulting pTOPO-0092 plasmid was sequenced to confirm that no PCR-derived mutations were introduced into the clone, and then was digested with NdeI and XhoI and the approximately ~1.5 kb insert corresponding to the *PSPTO_0092* CDS was ligated into the pET21a vector cut with the same enzymes to generate pET21a-0092. The pET21a-0092 plasmid was transformed into *E. coli* BL21(DE3). The same strategy was used to generate pET21a-0728, pET21a-2673 (AldB), pET21a-3064, pET21a-3323 and pET21a-3364 (AldC) (see S3 and S4 Tables for primers and strains).

For *E. coli* expression assays to monitor IAA production, the *E. coli* strains carrying the pET21a-DC3000 putative aldehyde dehydrogenase (Ald) constructs were grown in triplicate cultures overnight in LB media containing Amp with shaking at 37 °C. Overnight cultures were diluted 1/100 and incubated with shaking until an OD_600nm_ 0.4-0.6 was reached. Cultures were induced with IPTG (1 mM final concentration), supplemented with 0.25 mM IAAld and incubated with shaking for an additional 24 hrs. One mL samples were taken 1.5 hrs after IPTG induction to verify induction of the putative Ald proteins. This was done by centrifuging the samples, boiling the resulting cell pellets in SDS-PAGE buffer and loading equal amounts of cell lysate on an acrylamide gel for visualization of protein. Additional 1mL samples were taken at 24 hrs after IPTG induction, centrifuged to pellet cells and the resulting supernatants were frozen in liquid nitrogen and stored at -80 °C. The samples were analyzed for IAA production by LC-MS/MS [54].

### Protein expression and purification

The pET28a-AldA, pET28a-AldB, and pET28a-AldC constructs used to express protein for biochemical experiments were generated using NdeI and XhoI enzyme sites and transformed into *E. coli* BL21 (DE3) cells (Agilent Technologies). Cells were grown at 37 °C in Terrific broth containing 50 μg mL^-1^ Kan until OD_600nm_ = 0.8 and induced with 1 mM IPTG at 18 °C. Cells were harvested by centrifugation (4,500 × g; 15 min) and re-suspended in lysis buffer (50 mM Tris, pH 8.0, 500 mM NaCl, 25 mM imidazole, 10% glycerol, and 1% Tween-20). After sonication and centrifugation (11,000 x g; 30 min), the supernatant was loaded onto a Ni^2+^-NTA column (Qiagen) previously equilibrated with lysis buffer. Wash buffer (lysis buffer without Tween-20) was used to remove unbound proteins, and then bound Ald protein was eluted using wash buffer containing 250 mM imidazole. The His-tagged Ald protein was loaded onto a Superdex-200 26/60 size-exclusion column (GE healthcare) equilibrated in 25 mM Hepes (pH 7.5) and 100 mM NaCl. Fractions with Ald protein were pooled, concentrated to 10 mg mL^-1^, and stored at -80 °C. Protein concentrations were determined using molar extinction coefficients at A_280nm_ for each Ald, as calculated using ProtParam.

### Enzyme assays

Enzymatic activity of each Ald was measured by monitoring NADH formation (ε_340_ = 6220 M^−1^ cm^−1^) at A_340nm_ on an Infinite M200 Pro plate reader (Tecan). Standard assay conditions for Ald were 100 mM Tris•HCl (pH 8.0), 100 mM KCl in 200 μL at 25 °C. For specific activity determinations, the following substrate concentrations were used: 1 mM IAAld and either 1 mM NAD^+^ or 1 mM NADP^+^. For determination of steady-state kinetic parameters, reactions were performed in standard assay conditions with either fixed NAD^+^ (1.0 mM) and varied IAAld (0.05-2.5 mM) or with fixed IAAld (1.0 mM) and varied NAD^+^ (0.05-2.5 μM). All data were fit to the Michaelis-Menten equation, *v* = (*k*_cat_[S])/(*K*_m_ + [S]), using SigmaPlot.

### Protein crystallography and homology modeling

Crystallization of AldA was performed at room temperature using the vapor diffusion method in hanging drops of a 1:1 mixture of protein (10 mg mL^-1^) and crystallization buffer. Crystals of the AldA apoenzyme were obtained in 10% (w/v) PEG-8000, 100 mM HEPES, pH 7.5, and 8% (v/v) ethylene glycol. Crystals of the AldA•NAD^+^ and AldA•NAD^+^•IAA complexes were obtained in 8% (w/v) PEG-8000 and 100 mM Tris•HCl (pH 8.5) supplemented with either 5 mM NAD^+^ or 5 mM NAD^+^ and 5 mM IAA, respectively. Crystals were stabilized in cryoprotectant (crystallization solution with either 30% glycerol or 30% ethylene glycol) before flash freezing in liquid nitrogen for data collection at 100 K. Diffraction images were collected at beamline 19ID of the Advanced Photon Source at the Argonne National Lab. Diffraction data were indexed, integrated and scaled using HKL3000 [55]. The structure of AldA in complex with NAD^+^ was were solved by molecular replacement using PHASER [56] with betaine aldehyde dehydrogenase from *Staphylococcus aureus*, which shares 40% amino acid identity with AldA, as a search model (PDB: 4MPB; [57]. For iterative rounds of manual model building and refinement, COOT [58] and PHENIX [59] were used, respectively. The resulting model of AldA was used to solve the structures of the apoenzyme and NAD^+^•IAA complex by molecular replacement with PHASER. Model building and refinement was as described above. Data collection and refinement statistics are summarized in S2 Table. Atomic coordinates and structure factors were deposited in the RCSB Protein Data Bank (www.rcsb.org) as follows: AldA (5IUU); AldA•NAD^+^ (5IUV); and AldA•NAD^+^•IAA (5IUW).

### Homology modeling and computational docking

Molecular homology models of AldB and AldC were generated using the homology-modeling server of SWISS-MODEL with the 1.93 Å resolution crystal structure of AldA• NAD^+^•IAA (chain B) as a template. Molecular docking experiments were performed by Autodock vina (Version 1.1.2) [60]with standard protocols. Docking of IAAld (substrate) into the AldA active site used a 30 × 30 × 30 Å grid box with the level of exhaustiveness = 20. The position of NAD^+^ was fixed based on its position in the AldA•NAD^+^•IAA structure. Docking of IAAld yielded a calculated affinity of -5.9 to -4.8 kcal mol^-1^.

### Construction of *P. syringae ald* gene plasmid disruption mutants

To generate the *aldA::*pJP5603 insertion disruption strain, an ~0.5 kb SacI-XbaI genomic fragment internal to the *aldA (PSPTO_0092)* CDS was amplified from *P. syringae* DC3000 genomic DNA with the primers 0092SacIF and 0092XbaIR (see S4 Table for primer sequences). The resulting PCR fragment was cloned into the pBlunt II-TOPO vector (Invitrogen), transformed into *E. coli DH5α* and plated on LB media containing Kan. Several pTOPO-0092int clones were sequenced to verify that there were no PCR-derived mutations. The genomic fragment was then cloned into the pJP5603 KanR suicide vector [53] by digesting the pTOPO-0092int clone with *SacI* and *XbaI* and ligating the resulting genomic fragment into pJP5063 digested with SacI and XbaI to generate pJP5603-0092int. The pJP5603-0092int plasmid was transformed into *E.coli* DH5α *λpir* and introduced into *P. syringae* DC3000 via bacterial conjugation using the helper strain MM294A(pRK2013) (S3 Table) [61]. DC3000 trans-conjugates were selected for Rif^r^ and Kan^r^ resistance on NYG media containing Rif and Kan at 28 °C. The same strategy was used to generate *aldB::*pJP5603 and *aldC::*pJP5603 single mutants, as well as *aldA::*pJP5603 *aldB::*pJP5603-Tet, double mutant strains. To generate double mutants, a Tet^R^ version of the pJP5603-*aldB* insertion disruption suicide plasmid was used (see S3 and S4 Tables for primers and strains).

Plasmid disruption of *aldA* by pJP5603 was confirmed by PCR using primers M13F, 0092seqF, and 0092seqR. Disruption of the wild-type genomic copy was verified by amplification of an ~1.1 kb fragment with M13F and 0092seqF primers in the *aldA*:pJP56023 strain and the absence of a band of this size in wild-type DC3000 and *aldB*::pJP5603 and *aldC*::pJP5603 strains (S4 Fig. C and D). The same strategy was used to confirm all of the additional single and double *ald* mutants (see S3 and S4 Tables for strains and primers).

To generate the *aldA* complementing clone, p*AldA*, the *aldA* coding sequence and 5’ regulatory region were amplified from genomic DNA using primers 0092XhoIF and 0092EcoRIR. The resulting ~2 kb PCR product was cloned into the pBlunt II-TOPO vector (Invitrogen) to generate pTOPO-0092comp. This plasmid was then digested with XhoI and EcoRI and the 2 kb insert ligated into the broad host range plasmid pME6031 vector with Xho1 and EcoRI compatible ends to generate pME6031-0092 (p*AldA*) (S3 Table). The p*AldA* plasmid was introduced into the *aldA::*pJP5603 mutant strain via bacterial conjugation using the helper strain MM294A(pRK2013). DC3000 trans-conjugates were selected for Rif^r^, Kan^r^ and Tet^r^ resistance on NYG media containing Rif, Kan and Tet at 28 °C.

### Plant material and growth conditions

All *A. thaliana* transgenic lines and mutants used in this study were in the Col-0 background. The 35S:*YUC1* overexpression line [50] was obtained from Yunde Zhao. The *sid2*-*2* mutant [43] was obtained from Mary Wildermuth.

Plants were grown on soil in a growth chamber with a short-day photoperiod (8 h light/16 h dark) at 21°C and 75% relative humidity, with a light intensity of ~ 130 μEinsteins sec^-1^ m^-1^.

### *P. syringae* inoculation and quantification of bacterial growth

*A. thaliana* plants were infected at approximately four weeks of age. For surface inoculations plants were dipped into a solution containing *P. syringae* at approximately 3×10^8^ cells mL^-1^ (OD_600nm_ = 0.3), 10 mM MgCl_2_ and 0.02% (v/v) Silwet L-77 [62]. To quantify bacterial growth in the plant, whole leaves were sampled at various time points after inoculation, weighed to determine leaf mass, ground in 10 mM MgCl_2_ and then plated in serial dilutions on NYG media with rifampicin. Between four and six leaves were sampled per treatment, depending on the experiment. On the day of inoculation, leaves were sampled at 2 h after inoculation, surface sterilized with 15% (v/v) H_2_O_2_ and washed three times with sterile water before grinding to remove bacteria from the surface of the leaf. For syringe infiltrations, a solution containing 10^4^–10^5^ cells mL^-1^ (OD_600nm_ = 10^−5^-10^−4^) in 10 mM MgCl_2_ was injected into leaves using a 1-mL needleless syringe. Bacterial growth was monitored as described for dip inoculations, with the exception that leaves sampled on the day of inoculation were not subject to surface sterilization.

Quantification of disease symptoms following dip inoculation was carried out four days post inoculation. Leaves were categorized based on the presence and amount of chlorosis or yellowing of the leaf. For ~ 10 plants per each treatment, each leaf was individually assessed for percent of the leaf exhibiting chlorosis, ranging from leaves with no yellowing to leaves displaying >75% chlorosis.

## Acknowledgements

We thank A. Mutka, L. Strader, P. Levin, E. Mellgren and B. San Francisco for helpful discussion and comments on the manuscript. We also thank Andrew Mutka for providing plasmid pJP5603-Tet and the Proteomics and Mass Spectrometry Facility of the Donald Danforth Plant Science Center (St. Louis, MO) for hormone analysis.

## References

1. Jones JDG, Dangl JL. The plant immune system. Nature. 2006;444:323-329.

2. Xin XF, He SY. *Pseudomonas syringae* pv. tomato DC3000: a model pathogen for probing disease susceptibility and hormone signaling in plants. Annu Rev Phytopathol. 2013;51:473-498.

3. Chisholm ST, Coaker G, Day B, Staskawicz BJ. Host-microbe interactions: shaping the evolution of the plant immune response. Cell. 2006;124(4):803-814.

4. Preston GM. *Pseudomonas syringae* pv. *tomato*: The right pathogen, of the right plant, at the right time. Mol Plant Pathol. 2000;1(5):263-275.

5. Cuppels DA. Generation and Characterization of Tn5 Insertion Mutations in *Pseudomonas syringae* pv. tomato. Appl Environ Microbiol. 1986;52:323-327.

6. Staswick PE. JAZing up jasmonate signaling. Trends Plant Sci. 2008;13(2):66-71.

7. Katsir L, Schilmiller AL, Staswick PE, He SY, Howe GA. COI1 is a critical component of a receptor for jasmonate and the bacterial virulence factor coronatine. Proc Nat Acad Sci U S A. 2008;105(19):7100-7105.

8. Brooks DM, Hernández-Guzmán G, Kloek AP, Alarcón-Chaidez F, Sreedharan A, Rangaswamy V, et al. Identification and characterization of a well-defined series of coronatine biosynthetic mutants of *Pseudomonas syringae* pv. *tomato* strain DC3000. Mol Plant-Microbe Interact. 2004;17:162-174.

9. Brooks DM, Bender CL, Kunkel BN. The *Pseudomonas syringae* phytotoxin coronatine promotes virulence by overcoming salicylic acid-dependent defences in *Arabidopsis thaliana*. Mol Plant Pathol. 2005;6:629-639.

10. Uppalapati SR, Ishiga Y, Wangdi T, Kunkel BN, Anand A, Mysore KS, et al. The phytotoxin coronatine contributes to pathogen fitness and is required for suppression of salicylic acid accumulation in tomato inoculated with *Pseudomonas syringae* pv. tomato DC3000. Mol Plant-Microbe Interact. 2007;20:955-965.

11. Spaepen S, Vanderleyden J. Auxin and Plant-Microbe Interactions. Cold Spring Harb Perspect Biol. 2011(Apr; 3(4): a001438.).

12. Duca D, Lorv J, Patten CL, Rose D, Glick BR. Indole-3-acetic acid in plant-microbe interactions. Antonie Van Leeuwenhoek. 2014;106(1):85-125.

13. Patten CL, Blakney AJ, Coulson TJ. Activity, distribution and function of indole-3-acetic acid biosynthetic pathways in bacteria. Crit Rev Microbiol. 2013;39(4):395-415.

14. White FF, Ziegler S. Cloning of the genes for indoleacetic acid synthesis from *Pseudomonas syringae* pv. syringae. Mol Plant-Microbe Interact. 1991;4(207-210).

15. Koga J, Adachi T, Hidaka H. Purification and characterization of indolepyruvate decarboxylase. A novel enzyme for indole-3-acetic acid biosynthesis in *Enterobacter cloacae*. J Biol Chem. 1992;267(22):15823-15828.

16. Spaepen S, Versees W, Gocke D, Pohl M, Steyaert J, Vanderleyden J. Characterization of phenylpyruvate decarboxylase, involved in auxin production of *Azospirillum brasilense*. J Bacteriol. 2007;189(21):7626-7633.

17. Brandl MT, Lindow SE. Cloning and characterization of a locus encoding an indolepyruvate decarboxylase involved in indole-3-acetic acid synthesis in *Erwinia herbicola*. Appl Environ Microbiol. 1996;62(11):4121-4128.

18. Manulis S, Haviv-Chesner A, Brandl MT, Lindow SE, Barash I. Differential involvement of indole-3-acetic acid biosynthetic pathways in pathogenicity and epiphytic fitness of *Erwinia herbicola* pv. *gypsophilae*. Mol Plant Microbe Interact. 1998;11(7):634-642.

19. Aragon IM, Perez-Martinez I, Moreno-Perez A, Cerezo M, Ramos C. New insights into the role of indole-3-acetic acid in the virulence of *Pseudomonas savastanoi* pv. *savastanoi*. FEMS Microbiol Lett. 2014;356:184-192.

20. Woodward AW, Bartel B. Auxin: regulation, action, and interaction. Ann Bot (Lond). 2005;95(5):707-735.

21. Zhao Y. Auxin biosynthesis and its role in plant development. Ann Rev Plant Biol. 2010;61:49-64.

22. Enders TA, Strader LC. Auxin activity: Past, present, and future. Am J Bot. 2015;102(2):180-196.

23. Dobbelaere S, Croonenborghs A, Thys A, Vande Broek A, Vanderleyden J. Phytostimulatory effect of *Azospirillum brasilense* wild type and mutant strains altered in IAA production on wheat. Plant and Soil. 1999;212(2):153-162.

24. Jameson PE. Cytokinins and auxins in plant-pathogen interactions - An overview. Plant Growth Reg. 2000;32(2/3): 12.

25. Comai L, Kosuge T. Involvement of plasmid deoxyribonucleic acid in indoleacetic acid synthesis in *Pseudomonas savastanoi*. J Bacteriol. 1980;143(2):950-957.

26. Chen Z, Agnew JL, Cohen JD, He P, Shan L, Sheen J, et al. *Pseudomonas syringae* type III effector AvrRpt2 alters *Arabidopsis thaliana* auxin physiology. Proc Nat Acad Sci USA. 2007;104(50):20131-20136.

27. Wang D, Pajerowska-Mukhtar K, Culler AH, Dong X. Salicylic Acid Inhibits Pathogen Growth in Plants through Repression of the Auxin Signaling Pathway. Curr Biol. 2007;17(20):1784-1790.

28. Gonzalez-Lamothe R, El Oirdi M, Brisson N, Bouarab K. The conjugated auxin indole-3-acetic acid-aspartic acid promotes plant disease development. Plant Cell. 2012;24(2):762-777.

29. Mutka AM, Fawley S, Tsao T, Kunkel BN. Auxin promotes susceptibility to *Pseudomonas syringae* via a mechanism independent of suppression of salicylic acid-mediated defenses. Plant J. 2013;74:746–754.

30. Navarro L, Dunoyer P, Jay F, Arnold B, Dharmasiri N, Estelle M, et al. A plant miRNA contributes to antibacterial resistance by repressing auxin signaling. Science. 2006;312(5772):436-439.

31. Glickmann E, Gardan L, Jacquet S, Hussain S, Elasri M, Petit A, et al. Auxin production is a common feature of most pathovars of *Pseudomonas syringae*. Mol Plant Microbe In. 1998;11(2):156-162.

32. Rico A, Preston GM. *Pseudomonas syringae* pv tomato DC3000 Uses Constitutive and Apoplast-Induced Nutrient Assimilation Pathways to Catabolize Nutrients That Are Abundant in the Tomato Apoplast. Mol Plant-Microbe Interact. 1998;21(2):14.

33. Buell CR, Joardar V, Lindeberg M, Selengut J, Paulsen IT. The complete genome sequence of the Arabidopsis and tomato pathogen *Pseudomonas syringae* pv. *tomato DC3000*. Proc Natl Acad Sci U S A. 2003;100:10181-10186.

34. Bentley JA, Farrar KR, Housley S, Smith GF, Taylor WC. Some chemical and physiological properties of 3-indolylpyruvic acid. Biochem J. 1956;64(1):44-49.

35. Oberhansli T, Dfago G, Haas D. Indole-3-acetic acid (IAA) synthesis in the biocontrol strain CHA0 of *Pseudomonas fluorescens*: role of tryptophan side chain oxidase. J Gen Microbiol. 1991;137(10):2273-2279.

36. Spaepen S, Vanderleyden J, Remans R. Indole-3-acetic acid in microbial and microorganism-plant signaling. FEMS Microbiol Rev. 2007;31(4):425-448.

37. Xie B, Xu K, Zhao HX, Chen SF. Isolation of transposon mutants from *Azospirillum brasilense* Yu62 and characterization of genes involved in indole-3-acetic acid biosynthesis. FEMS Microbiol Lett. 2005;248(1):57-63.

38. Singh S, Brocker C, Koppaka V, Chen Y, Jackson BC, Matsumoto A, et al. Aldehyde dehydrogenases in cellular responses to oxidative/electrophilic stress. Free Radical Biol & Med. 2013;56:89-101.

39. Weiner H, Wang X. Aldehyde dehydrogenase and acetaldehyde metabolism. Alcohol and Alcoholism. 1994;2:141-145.

40. Steinmetz CG, Xie P, Weiner H, Hurley TD. Structure of mitochondrial aldehyde dehydrogenase: the genetic component of ethanol aversion. Structure. 1997;5(5):701-711.

41. Koncitikova R, Vigouroux A, Kopecna M, Andree T, Bartos J, Sebela M, et al. Role and structural characterization of plant aldehyde dehydrogenases from family 2 and family 7. Biochem J. 2015;468(1):109-123.

42. Robert-Seilaniantz A, Grant M, Jones JD. Hormone crosstalk in plant disease and defense: more than just jasmonate-salicylate antagonism. Annu Rev Phytopathol. 2011;49:317-343.

43. Wildermuth MC, Dewdney J, Wu G, Ausubel FM. Isochorismate synthase is required to synthesize salicylic acid for plant defense. Nature. 2001;414(6863):562-565.

44. Brocker C, Vasiliou M, Carpenter S, Carpenter C, Zhang Y, Wang X, et al. Aldehyde dehydrogenase (ALDH) superfamily in plants: gene nomenclature and comparative genomics. Planta. 2013;237(1):189-210.

45. Milo R, Last RL. Achieving diversity in the face of constraints: lessons from metabolism. Science. 2012;336(6089):1663-1667.

46. Perez-Miller SJ, Hurley TD. Coenzyme isomerization is integral to catalysis in aldehyde dehydrogenase. Biochem. 2003;42(23):7100-7109.

47. Mashiguchi K, Tanaka K, Sakai T, Sugawara S, Kawaide H, Natsume M, et al. The main auxin biosynthesis pathway in Arabidopsis. Proc Natl Acad Sci U S A. 2011;108(45):18512-18517.

48. Ouyang J, Shao X, Li J. Indole-3-glycerol phosphate, a branchpoint of indole-3-acetic acid biosynthesis from the tryptophan biosynthetic pathway in *Arabidopsis thaliana*. Plant J. 2000;24(3):327-333.

49. Park JE, Park JY, Kim YS, Staswick PE, Jeon J, Yun J, et al. GH3-mediated auxin homeostasis links growth regulation with stress adaptation response in Arabidopsis. J Biol Chem. 2007;282(13):10036-10046.

50. Zhao Y, Christensen SK, Fankhauser C, Cashman JR, Cohen JD, Weigel D, et al. A role for flavin monooxygenase-like enzymes in auxin biosynthesis. Science. 2001;291(5502):306-309.

51. Daniels MJ, Barber CE, Turner PC, Sawczyc MK, Byrde RJW, Fielding AH. Cloning of genes involved in pathogenicity of *Xanthomonas campestris* pv. *campestris* using the broad host range cosmid pLAFR1. EMBO J. 1984;3:3323-3328.

52. Sreedharan A, Penaloza-Vazquez A, Kunkel BN, Bender CL. CorR regulates multiple components of virulence in *Pseudomonas syringae* pv. *tomato* DC3000. Mol Plant-Microbe Interact. 2006;19:768-779.

53. Penfold RJ, Pemberton JM. An improved suicide vector for construction of chromosomal insertion mutations in bacteria. Gene. 1992;118:145-146.

54. Chen Q, Zhang B, Hicks LM, Wang S, Jez JM. A liquid chromatography-tandem mass spectrometry-based assay for indole-3-acetic acid-amido synthetase. Anal Biochem. 2009;390(2):149-154.

55. Minor W, Cymborowski M, Otwinowski Z, Chruszcz M. HKL-3000: the integration of data reduction and structure solution‐‐from diffraction images to an initial model in minutes. Acta Crystallogr D Biol Crystallogr. 2006;62(Pt 8):859-866.

56. McCoy AJ, Grosse-Kunstleve RW, Adams PD, Winn MD, Storoni LC, Read RJ. Phaser crystallographic software. J Appl Crystallogr. 2007;40(Pt 4):658-674.

57. Chen C, Joo JC, Brown G, Stolnikova E, Halavaty AS, Savchenko A, et al. Structure-based mutational studies of substrate inhibition of betaine aldehyde dehydrogenase BetB from *Staphylococcus aureus*. Appl Env Microbiol. 2014;80(13):3992-4002.

58. Emsley P, Lohkamp B, Scott WG, Cowtan K. Features and development of Coot. Acta Crystallogr D Biol Crystallogr. 2010;66(Pt 4):486-501.

59. Adams PD, Afonine PV, Bunkoczi G, Chen VB, Davis IW, Echols N, et al. PHENIX: a comprehensive Python-based system for macromolecular structure solution. Acta Crystallogr D Biol Crystallogr. 2010;66(Pt 2):213-221.

60. Trott O, Olson AJ. AutoDock Vina: improving the speed and accuracy of docking with a new scoring function, efficient optimization, and multithreading. J Comput Chem. 2010;31(2):455-461.

61. Finan TM, Kunkel B, de Vos GF, Signer ER. Second symbiotic megaplasmid in *Rhizobium meliloti* carrying exopolysaccharide and thiamine synthesis genes. J Bacteriol. 1986;167(1):66-72.

62. Kunkel BN, Bent AF, Dahlbeck D, Innes RW, Staskawicz BJ. *RPS2*, an Arabidopsis disease resistance locus specifying recognition of *Pseudomonas syringae* strains expressing the avirulence gene *avrRpt2*. The Plant cell. 1993;5(8):865-875.

